# Early Spring Snowmelt and Summer Droughts Strongly Impair the Resilience of Key Microbial Communities in Subalpine Grassland Ecosystems

**DOI:** 10.1101/2021.03.15.435477

**Authors:** Farhan Hafeez, Lionel Bernard, Franck Poly, Jean-Christophe Clément, Thomas Pommier

**Author notes:** ^*^***Corresponding author*** : Thomas Pommier, *Laboratoire d’Ecologie Microbienne LEM, INRA UMR 1418, CNRS UMR 5557, Université de Lyon, F◻69622 Villeurbanne Cedex, France,.

## Abstract

Subalpine grassland ecosystems are important from biodiversity, agriculture, and touristic perspectives but their resilience to seasonally occurring climatic extremes is increasingly challenged with climate change, accelerating their vulnerability to tipping points. Microbial communities, which are central in ecosystem functioning, are usually considered as more resistant and highly resilient to such extreme events due to their functional redundancy and strong selection in residing habitats. To investigate this, we explored soil microbial responses upon recurrent summer droughts associated with early snowmelt in subalpine grasslands mesocosms set-up at the Lautaret Pass (French Alps). Potential respiration, nitrification and denitrification were monitored over a period of two growing seasons along with quantification of community gene abundances of total bacteria as well as (de)nitrifiers. Results revealed that droughts had a low and short-term adverse impact on bacterial total respiration supporting their hypothesized high resilience, i.e., resistance and ability to recover. Nitrification and abundances of the corresponding functional guilds showed relatively strong resistance to summer droughts but declined in response to early snowmelt. This triggered a cascading effect on denitrification but also on abundances of denitrifying communities which recovered from all climatic extremes except from the summer droughts where nitrifiers were collapsed. Denitrification and respective functional groups faced high impact of applied stresses with strong reduction in abundance and activity of this specialized community. Although, consequently lower microbial competition for nitrate may be positive for plant biomass production, warnings exist when considering the potential nitrogen leaching from these ecosystems as well as risks of greenhouses gases emission such as N_2_O.

## 1. Introduction

Subalpine grassland ecosystems are important for biodiversity, agriculture, and a wide range of key services (Pecher et al., 2017; Zoderer et al., 2016; Tappeiner et al., 2008; Grêt-Regamey et al., 2012). These grasslands are under extreme pressure of climatic changes drastically affecting their functioning and disrupting the linked ecosystem services, *e.g.,* plants productivity, nutrient cycling etc. (Berauer et al., 2019; Bernard et al., 2019). Considering the continuous human perturbations and global changes (Berauer et al., 2019), it is direly required to investigate whether these grasslands are highly vulnerable, have developed resilience (comprising of resistance: ability to maintain the existing state, and recovery characteristics: to recover from disturbances) (Ingrisch and Bahn, 2018; Hodgson et al., 2015) or have attained an altered recovery state (Seabloom et al., 2020) over time which is crucial in conserving ecosystems functions (Donohue et al., 2016).

The multiple effects of global climatic changes not only impact biodiversity and ecosystem services (Berauer et al., 2019; Schirpke et al., 2013, Engler et al., 2011) but also their associated resilience including both resistance (Bernard et al., 2019; Wipf and Rixen, 2010) and recovery in Alpine grasslands (Ingrisch et al., 2020; Karlowsky et al., 2018). The effect of recurring climatic extremes on microbial functioning in these grasslands needs to be assessed as they may shift the fate of particular biogeochemical functions especially when resilience limits are crossed and tipping points (where a potentially irreversible change is about to occur or is already occurring *sensu*, Schnellnhuber et al. 2006) are reached. Such estimations are crucial to understand and possibly protect the stability of ecosystems and inhabiting species (de Vries and Shade, 2013; Griffiths and Philippot, 2013), but also to forecast unforeseen consequences of adverse climatic extremes. Where weather extremes including droughts are predicted to be severe (Fuchslueger et al., 2019; Karlowsky et al., 2018), Alpine grassland ecosystems are expected to face less snow falls that ultimately lead to subsequent droughts (Klein et al., 2016; Gobiet et al., 2014). However, their vulnerability to collapse or support a particular function towards the combined (cumulative) impact of different weather extremes is poorly documented.

Microbial communities are crucial actors in ecosystem functioning (Chen et al., 2020; Bardgett and van der Putten, 2014) and respond strongly to environmental variability (Piton et al., 2020). These are critical actors in estimating the functional stability (de Vries and Shade 2013; Griffiths et al., 2000), and they are highly diverse and functionally redundant across a range of environments. Consequently, microbial communities are usually expected to be highly resilient to climatic fluctuations as compared to many other environmental actors albeit the strong selection under local harsh climatic conditions (Capdeville et al., 2019; Griffiths and Philippot, 2013, Benot et al., 2014). Despite these yet known inherent capabilities, whether these microbial communities can withstand to correlated acceleration of weather extremes is largely unknown. Especially, assessment of the functional boundaries of the subalpine grasslands and linked ecosystems services is critical since they are susceptible to extreme events (Bernard et al., 2019).

The characteristics of existing plant species and nutrient availability as well as their interaction with microorganisms to ultimately respond to climatic changes are important in assessing soil functioning resilience (Bernard et al., 2020). For example, any shift in native plant community can influence the soil nutrient usage leading to selection in the pool of residing microbes hence the corresponding functions (Legay et al., 2016; de Vries et al., 2012). Similarly, weather extremes directly influencing plant community may also impact microbes (Bardgett et al., 2013; de Vries and Shade, 2013, Saccone et al., 2013). Earlier, summer droughts in subalpine grasslands were reported to increase leaf senescence while reducing plant productivity (Benot et al., 2014), as well as affecting N cycling-related microbial community and functions (Dai et al., 2020; Cantarel et al., 2012).

Here we determined the short-term effect of drought, snow removal, and their additive (recurrent) effect on microbial activities and size of involved functional guilds and following responses, to measure the resistance and recovery, respectively. Since microbes, driving broad (*e.g.,* respiration, decomposition) and specialized soil processes (*e.g.* (de)nitrification), may respond differentially to various perturbations (Dai et al., 2020; Jusselme et al., 2016; Chaer et al., 2009; Wertz et al., 2007), we conducted our work both at broad and functional scales. Investigating the response of these grasslands is crucial since the variable impact of natural and manmade perturbations might have considerably influenced nutrient cycling including that of N (Galloway et al., 2004), but also the involved microbial functional guilds. First, we tested how total bacterial community (16S *rRNA*) and related substrate-induced respiration (SIR) were influenced under droughts, snow removal and their cumulative impact and how much they recovered from these stresses. Then we investigated nitrogen (N) cycling (that is critical in ecosystem sustainability (Chen et al., 2020; Kuypers et al., 2018)) by determining the abundances (nitrifiers (*nxrA*, *NS*), denitrifiers (*nirK*, *nirS*, *nosZ*)) and activities (nitrification and denitrification enzyme activities -NEA and DEA, respectively) of involved microbial communities, since they carry implications for plant productivity (loss of N - an important plant nutrient) and environment (N_2_O emissions-global warming) (Dai et al., 2020; Galloway et al., 2004).

The current work was carried out on functionally contrasted and assembled grasslands set up in a common garden at Lautaret Pass with various sampling campaigns during two growing seasons. Based on an assumed functional redundancy (Jia and Whalen 2020; Griffiths and Philippot 2013), we hypothesized that broad scale function i.e., SIR and relevant microbial community should resist and/or recover once the stresses/extreme events are over. SIR being representative of total microbial communities is expected to be influenced by soil moisture and abundance of *16S rRNA* (Hallin et al., 2009). However, more specific functions of N-cycling should be more sensitive because we expect comparatively less functional redundancy and, despite being able to resist to single stress, the combined effects (SR+D) may trigger a tipping point, disturbing various steps of these processes. Moreover, any shift in soil moisture, substrate availability and abundance of denitrifiers can strongly impact DEA through affecting micro-environment conditions (Philippot et al., 2007) both under drought and no drought, while NEA might be sensitive to ammonium availability (Jia and Conrad 2009) and snow removal. To address these questions, we quantified the immediate responses of microbial communities’ abundances and processes to droughts and early snowmelt extreme events to explore the resistance and the following recovery across subalpine grassland ecosystems.

## 2. Materials and methods

### 2.1. Experimental site, Mesocosms and Treatments’ implementation

The study was set up at Lautaret pass proximity in southeast France (2,100 m a.s.l, 45°02’13’’N, 6°24’01’’E) with a multifactorial experimental design where 128 mesocosms (n=128), 50-cm width and 40-cm depth, were installed in a common garden beside subalpine grasslands. Mesocosms were filled with previously sieved (5-mm) *in-situ* soil and planted with varying relative abundances of four species ranging along a plant economics spectrum. Species richness and functional composition remained stable throughout the whole experimentation due to low plant mortality. No snow removal (NSR) and snow removal (SR) treatments (n=64+64) were performed and for each of the NSR and SR there was an additional (sub) climate change treatment, i.e., maintained average soil moisture (ND) or drought (D) (n=32+32). Treatments receiving no snow removal with maintained average soil moisture were named as control (CTRL). All the 128 Mesocosms were installed inside 8 blocks. Within each of the 8 blocks with or without snow removal, there were two sub-blocks either maintained at long-term average soil moisture or subjected to summer drought. Within each sub-block 8 mesocosms were installed and 4 were submitted to plant cuttings and fertilization (CF), while the other half remained untreated (NCF) (n=16+16). During spring 2014, slow-release fertilizer was applied as pellets in the CF mesocosms (150 kg N/ha). To simulate early snowmelt, snow cover was removed one month earlier than the average snowmelt period which ultimately simulated a shorter snow-covered period followed by an early spring drought. The drought treatment was implemented using rainout shelters during spring, summer and fall. These shelters were thus in use during the whole vegetation season and all mesocosms were watered manually to maintain long-term average soil moisture. This manual watering was discontinued to achieve drought that reduced soil moisture for more than one month in August - September 2013 and June - July 2014.

### 2.2. Soil sampling campaigns

Five sampling campaigns in total were carried out at different time intervals including June 2013 (T0), September 2013 (T1), May 2014 (T2), July 2014 (T3) and September 2014 (T4). 1^st^ and 2^nd^ D were applied in August - September 2013 and June - July 2014 respectively (Fig. 1). The SR treatment was applied before T2 sampling campaign. Resistance to 1^st^ D of the microbial communities was measured at T1, resistance to 1^st^ SR and recovery from 1^st^ D at T2, resistance to 2^nd^ D and recovery from 1^st^ SR at T3; and recovery from overall D and SR treatments was evaluated at T4 sampling campaign (Fig. 1). No difference at T0 observed since no treatment was applied, and control was already placed at each date - T1 to T4, so we choose to compare only these on graph. During all sampling campaigns composite soil samples (5 cores mixed together, 0-10 cm depth) were collected from all 128 mesocosms (128×5: n=640). Fresh soil samples were sieved at 2-mm, homogenized, and brought to the laboratory to measure the enzyme activities in fresh soil while a sub-sample was stored at −20°C for DNA extraction. Soil properties including soil moisture content, SOM% (soil organic matter), soil pH, soil %C, soil %N, total dissolved nitrogen-TDN, and the dissolved organic nitrogen-DON, soil NH_4_^+^, and soil NO_3_^−^ were also measured in an associated work with focus on the effects of local environmental changes on subalpine grasslands functioning (Bernard 2017). Specifically, subsamples of 5 g fresh soil were dried at 70°C for 1 week to determine soil moisture content (in g.g-1 dw calculated as the 70°C dry soil weight relative to the fresh mass), followed by 4 h at 550° C to determine SOM (calculated as the 550°C soil weight relative to the 70°C dry mass). 10 mg soil subsample were air dried, ground to powder and analyzed for soil total C and N concentrations using a C/N elemental analyzer (FlashEA 1112, ThermoFisher Scientific, MA, USA). Soil pH was measured in a 1:4 (air dry soil/distilled water) solution. Soil N-NH_4_^+^ and N-NO_3_^−^, N-TDN and N-DON were determined in separated 10 mg fresh soil subsamples extracts (K_2_SO_4_, 0.5 M) using an FS-IV colorimetric chain (OI-Analytical, College Station, TX, USA) (Jones and Willett, 2006).

**Fig. 1.**
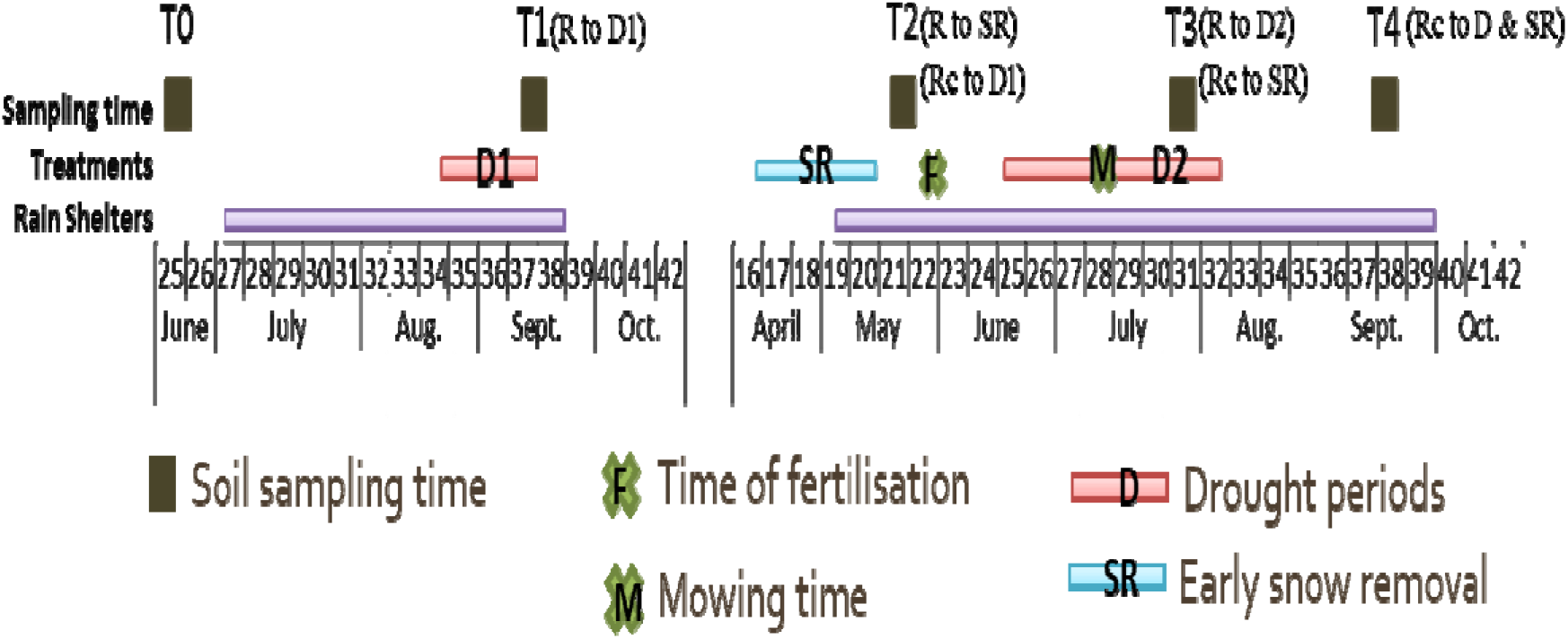
Experimental timeline showing the implementation of various treatments as well as the sampling dates. R and Rc represent resistance and recovery respectively.

### 2.3. Soil DNA extractions and quantifications

DNA was extracted from soil stored at −20°C using the PowerSoil® DNA Isolation Kit (Mo Bio Laboratories Inc. USA) and DNA concentration was determined by using the Quant-iTTM PicoGreen dsDNA Assay Kit (Invitrogen, Carlsbad, CA, USA).

### 2.4. Microbial community abundances

The abundance of the total bacterial communities was estimated in all DNA extracts using a *16S rRNA* primer-based quantitative PCR (qPCR) assays as described by Lane (1991). Final reaction volume was 20-μl, with 300-nM of forward and reverse primers; 1 X LightCycler DNA Master Mix SYBR® Green I (Roche Diagnostics); 1-ng of sample DNA or 10^8^ –10^2^ copies of the standard DNA pQuantAlb16S plasmid (Zouacheet al., 2012). The abundances of the *NS, nxrA, nirS, nirK and nosZ* genes were estimated using the primers developed previously (Attard et al., 2011; Henry et al., 2004; Henry et al., 2006; Kandeler et al., 2006). The total volume of the reaction was made 20 μl for *nxrA* and *nirK* each, with concentration of 500 nM for *nxrA* while 1μM for *nirK* for both forward as well as reverse primers; 10 μL of the SYBR® Green I kit was used for the *nxrA* and *nirK*, and 0.4 μg of T4 protein (Qbiogene, Carlsbad, CA USA) was used for *nirK*. The total volume of the reaction was made 25 μl for *NS*, *nirS* and *nosZ*, with concentration of 400 nM for *NS* while 1μM for *nirS* and *nosZ* both for forward as well as reverse primers; 12.5 μL of the SYBR® Green I kit was used for the *NS*, *nirS* and *nosZ*; for *nirS* the T4 protein was 0.5 μg while it was 0.4 μg for *nosZ*. The curves for the standards for *NS*, *nxrA*, *nirS, nosZ* and *nirK* were acquired after the quantitative PCR assays with regular dilutions of already known concentrations of the plasmid DNA with corresponding genes (10^7^ - 10^1^ number of gene copies). Quantification of all gene community abundances was conducted out on a lightcycler 480 (Roche Diagnostics). The amplification conditions used for qPCR assays were: 10 min at 95°C for each 40 cycles for *16S rRNA*, 45 for *nxrA*, *NS*, *nirK* and *nirS* and 40 for *nosZ* (after 6 cycles of touchdown); 95°C for 15 s for *16S rRNA*, *nirK*, *nirS*, *nosZ* and for 30 s for *nxrA* and NS; 63°C for 1 min for *16S rRNA*, 55°C for 45 s for *nxrA* and 66°C for 30 s for *NS*, 30 s for *nirK* and *nirS*, 60°C for 15 s for *nosZ*. 72°C for 30 s for 16S *rRNA*, *nirK*, *nirS* and *nosZ*, 45 s for *nxrA*, 1 min for NS; and additional step of 80°C for 15 s *nirS* and *nosZ*. Then 95°C for 5 s for 16S *rRNA* and *nxrA*, 1 s for *NS*, *nirK* and *nirS* and 15 s for *nosZ*; 65°C for 1 m for *16S rRNA* and *nxrA*, 68°C for 20 s for *NS* and *nirK*, 60°C for 20 s for *nirS* and 15 s for *nosZ*. And a continuous increase to 98°C for *16S rRNA*, *nxrA*, *NS, nirK* and *nirS* and 95°C for *nosZ* to determine the melting point; and finally, 10 s at 40°C for *16S rRNA*, *nxrA*, *NS*, *nirK* and *nirS* while 30 s for *nosZ* for cooling. At least two independent quantitative PCRs were done for every sample while the average was used. Co-amplification of the standards and the dilution of samples were used to control any inhibition in the quantification.

### 2.5. Microbial activities

Susbtrate Induced Respiration MicroResp. (SIR) was measured as CO_2_ production using MicroResp plates (Chapman et al., 2007). Soil was added to the wells of 96-well plates. 0.5-ml of a nutritive solution including glucose (1.2-mg of C-glucose g of dried soil) was added then incubated at 28°C. CO_2_ concentrations were measured using a TECAN spectrophotometer after 6-hours of incubation and the slope of the linear regression was used to estimate aerobic respiration as the CO_2_ produced (ppmV CO_2_ h^−1^ g^−1^).

Nitrifying enzyme activity (NEA) was measured on fresh soil samples as described by Patra et al. (2005). In brief, the fresh soil sample (equal to 3-g dry weight) was incubated with 3 ml solution of N-NH_4_ (200 μg N-(NH_4_)_2_SO_4_ g^−1^) in the flask for 72-hours at 180-rpm and 28°C. The flask was sealed with parafilm® (Pechiney Plastic Packaging, Menasha, WI, USA) to produce the aerobic conditions. With distilled water, the total volume of the suspension was made up to 15-ml. During the incubation, at regular intervals of 5, 24, 48 and 72-hours soil slurry was filtered with a pore size of 0.2-μm. The rates of production of NO_2_/NO_3_ were measured by ionic chromatography (DX120. Dionex, Salt Lake City, UT, USA).

Denitrification enzyme activities (DEA) was measured by incubating the fresh soil sample in a 150-ml airtight plasma-flask sealed by a rubber stopper for 4-hours at 28°C followed by the N_2_O measured each 30 minutes for 3-hours a gas chromatograph coupled to a micro-catharometer detector (lGC-R3000; SRA instruments) using the method adapted from Dassonville et al. (2011). Briefly, a substrate solution (2-ml) containing glucose (0.5-mg of C-glucose g^−1^ of dried soil), glutamic acid (0.5-mg of C-glutamic acid g^−1^ of dried soil) and potassium nitrate (50-μg of N-KNO_3_ g^−1^ of dried soil) were added to the soil. A 90:10 He-C_2_H_2_ atmosphere provided anaerobic conditions and inhibited N_2_O-reductase activity using the acetylene inhibition technique that blocks the last step in the denitrification pathway resulting in the accumulation of N_2_O (Yoshinari et al., 1977). The soil was incubated for 4-hours at 28°C and after incubation the amount of N_2_O was measured each 30 minutes for 3-hours. The amount of N_2_O produced was measured using a gas chromatograph coupled to a micro-catharometer detector (lGC-R3000; SRA instruments).

### 2.6. Statistics and Structural Equation Model

The data obtained were analyzed using JMP8^®^ (SAS Institute Inc., SAS Campus Drive, NC, USA). First, we did t-test to know the distribution of data (which was mostly not normally distributed except for SIR) and then ANOVA (for normally distributed data) and Kruskal-Wallis pairwise test (suited for non-normally distributed data) were applied to know which treatment is different from the other for studied parameters at various sampling times. Similarly, Kruskal-Wallis pairwise comparison was carried out also for the data which remained non-normally distributed even after a log transformation. In addition to linear modeling, for block effect, and repeated measures considerations (to consider time component or sampling from same place at different times), we used Cluster analyses approach using R (the package of R for hierarchical cluster analysis is ‘cluster’ and the function is hclust and to make dendogram we used function dendogram) to determine the effect of treatments within each block (to investigate the block effect) at different sampling times (to investigate the repeated measures) on each other. Structural Equation Model (SEM) was constructed using AMOS (Amos Development Corporation, Crawfordville, Florida, USA) to link the environmental and microbiological variables to investigate the relationship between these parameters under different climatic extremes. The fitness of the model was assessed by using the χ^2^ test and the model was considered as correct if the χ^2^ test produced a non-significant p-value - p>0.05. Since the factors controlling the microbial functioning under the drought and/or snow removal treatments were changing upon these treatments, the hypothesized models were changed to increase the level of significance and to fit the model. SEM were constructed to understand the relationship and path coefficients among the various causal environmental and microbial factors under D, SR, SR+D treatments. The choices for SEMs were made considering the parameters showing strong correlations as obtained through analyses of linear regression modeling. Initial SEMs were constructed for each sampling interval at each period of time. Then the strongly correlated parameters were linked through SEMs to understand the variable impact of treatments for different parameters hence variability was also considered.

## 3. Results

### 3.1. Changes in Broad Scale Function and Community

Soil moisture in CTRL showed the significant effect of the D, SR and SR+D treatments at all sampling times (Fig. 2a; Fig. S1). The resistance and recovery of the N-related microbial activities both for nitrification and denitrification and the abundances of corresponding genes were calculated as the relative change in D, SR, and SR+D treatments in comparision to CTRL. SIR was found to be significantly (p=0.001) lowered upon both droughts and was recorded up to 89% of CTRL (43.83 ppmV CO_2_ h^−1^ g^−1^ ± 0.58) but recovered ultimately (p=0.639). Although the interaction of SR+D treatments marginally (p=0.0123) affected the SIR, it was not affected by SR alone (p=0.625). It was noteworthy that the abundance of the total bacterial community (as expressed by 16S *rRNA* gene copies) followed a similar trend as SIR and was significantly (p=0.0006) reduced up to 56.3% of CTRL (4.87 × 10^8^ copies per g of dry soil) with a complete recovery observed at T4 (Fig. 2; Fig. S1).

**Fig. 2.**
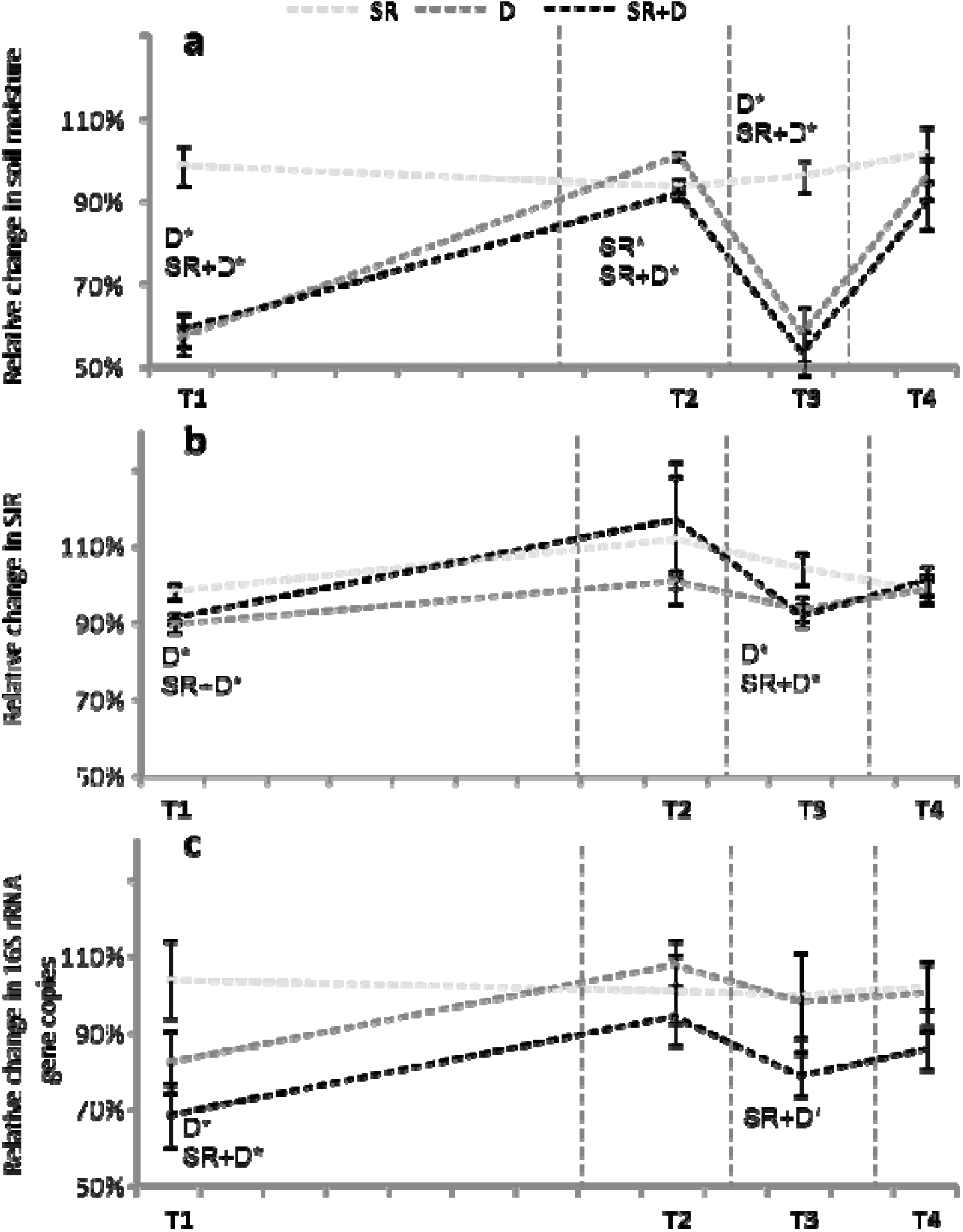
Relative change compared to CTRL for different treatments at various sampling dates **a)** in soil moisture; **b)** in SIR-MicroResp™; **c)** in abundance of total bacterial community (number of 16S *rRNA* gene copies g^−1^ dry soil); Light grey dotted lines show variations in time of the variables in the snow removal (SR) treatment relative to CTRL; dark grey dotted lines show variations of the Drought (D) treatment relative to CTRL; and the black dotted lines show the variations of cumulative SR+D treatment relative to CTRL.

### 3.2. Response of N Cycling Processes to Droughts and Snowmelt

Nitrification (NEA) showed resistance to both droughts at T1 and T3 and thus remained independent of the change in moisture content. Whereas, there was a significant reduction in NEA (up to 76% of CTRL) as observed upon SR treatment at T2 (Fig. 3; Fig. S2). To assess linkages, multiple regression analyses were carried out for the parameters in addition to moisture contents and the findings showed that the NEA was positively correlated to NH_4_, %N, TDN and *nxrA* (r = 0.56, 0.20, 0.31, 0.20; p < 0.005). On the contrary, DEA was significantly affected by both droughts and was found to be lowered up to 85% and 68% of CTRL upon first and second drought, respectively (Fig. 4; Fig. S3). Recovery from the first drought was observed for DEA at T2 except for the interaction of SR+D treatment. However, the second drought (at T3, Fig. 4) strongly (p=0.0001) impacted DEA which did not recover from SR. Hence, at T4 there was an incomplete recovery in denitrification acitivity as the DEA for SR+D mesocosms could not return to the initial levels (0.362 μg N-N_2_O/g dry soil/h ± 0.022).

**Fig. 3.**
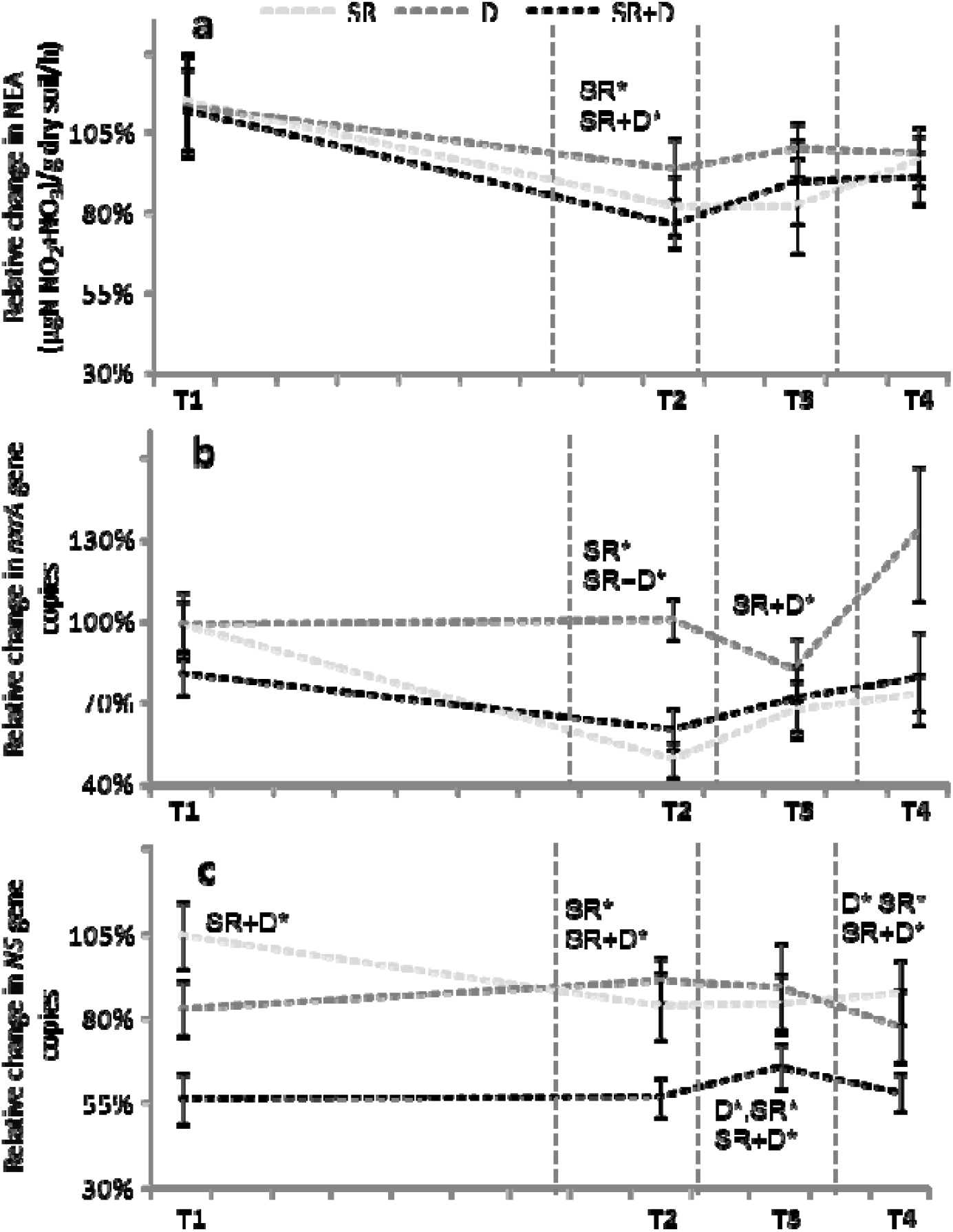
**a)** Relative change compared to CTRL in **a)** nitrification enzyme activities (μgN-(NO_2_+NO_3_)/g dry soil/h); **b)** in abundance of *nxrA* nitrite reductases (number of *nxrA* gene copies g^−1^ dry soil); **c)** in abundance of *NS* nitrite reductases (number of *NS* gene copies g^−1^ dry soil); Light grey dotted lines show variations in time of the variables in the snow removal (SR) treatment relative to CTRL; dark grey dotted lines show variations of the Drought (D) treatment relative to CTRL; and the black dotted lines show the variations of cumulative SR+D treatment relative to CTRL.

**Fig. 4.**
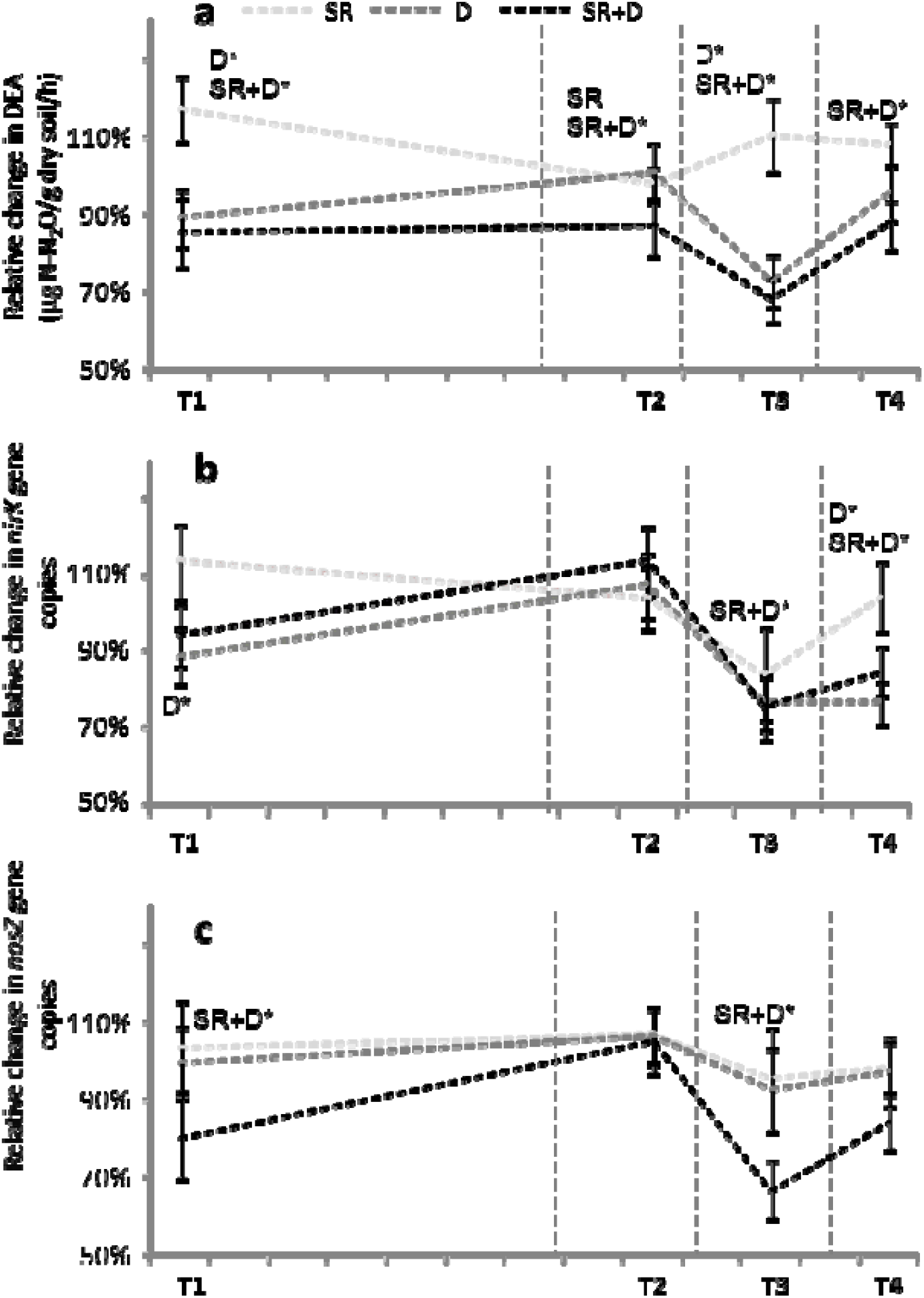
**a)** Relative change compared to CTRL in **a)** denitrification enzyme activities (μg N-N_2_O/g dry soil/h); **b)** in abundance of *nirK* denitrifiers (number of *nirK* gene copies g^−1^ dry soil); **c)** in abundance of *nosZ* denitrifiers (number of *nosZ* gene copies g^−1^ dry soil); Light grey dotted lines show variations in time of the variables in the snow removal (SR) treatment relative to CTRL; dark grey dotted lines show variations of the Drought (D) treatment relative to CTRL; and the black dotted lines show the variations of cumulative SR+D treatment relative to CTRL.

### 3.3. Impact on N Cycling Functional Guilds

Our results revealed that the abundances of N cycling functional guilds including nitrifiers and denitrifiers were strongly impacted by both D and SR climatic extremes (p=0.001). We observed that the *Nitrospira* and *Nitrobacter* (*nxrA* and *NS)* abundances were significantly (p=0.001) reduced upon applied stresses and could recover only for *nxrA* (Fig. 3). The denitrifiers were particularly impacted by the second drought, that strongly lowered their gene abundances. At T4, the community gene abundances of *nirS* and *nosZ* denitrifiers were completely recovered. However, *nirK* denitrifiers could not recover in agreement to the observations as recorded for DEA (Fig. 4; Fig. S3). The multiple correlation analyses as discussed in the next section showed the strong positive correlations of DEA with the %C, %N, DON, SOM, soil moisture, NEA as well as with the abundances of denitrifying genes. However, these correlations were found more prominent in the climate change treatments than the CTRL especially for soil moisture. For example, the Spearman correlation coefficient showed a non-significant correlation between soil moisture and *nirK* denitrifiers (r = 0.04; p > 0.652) only for the CTRL whereas it was positive in SR+D treatment (r = 0.33; p < 0.005).

### 3.4. Microbial Variables and Climatic Treatments, a Structural Equation Modeling

Fig. 5 represents how the various environmental (soil moisture, NH_4_, NO_3_ etc.) and microbial parameters (gene abundances) influenced the broad scale and N-related functional activities of microbial communities at different levels of drought and snow removal as well as by their interaction. The initial SEM was constructed for each sampling interval to observe how the various environmental and microbial factors influenced the broad scale (SIR) and N-related (NEA, DEA) functional microbial activities over each period of time. Afterwards, we separately analyzed the most correlated parameters from the results of the linear regression modeling and these parameters were linked to fit the SEM for drought and snow removal treatments but also for their interaction. The SEM revealed that the SIR was linked to soil moisture and *16S rRNA* for various treatments at different sampling times except at T1 and T2 (χ^2^ = 14.35, p = 0.35; Fig. 5). In contrast, NEA was never associated to soil moisture and was found to be mainly linked with soil NH_4_ concentration and community gene abundances of the *NS* nitrifiers, except for the SR treatment. Interestingly, under droughts, major part of the DEA variance (up to 55%) was explained by soil moisture with *nirS* as dominant denitrifiers except at T3 when it was *nosZ*. It was also observed that the overall impact of snow removal and drought was variable for different denitrifiers with severe impact on *nirK* which could not recover even by the end of the experiment. No significant effects of plant functional composition and fertilization on microbial parameters were recorded.

**Fig. 5.**
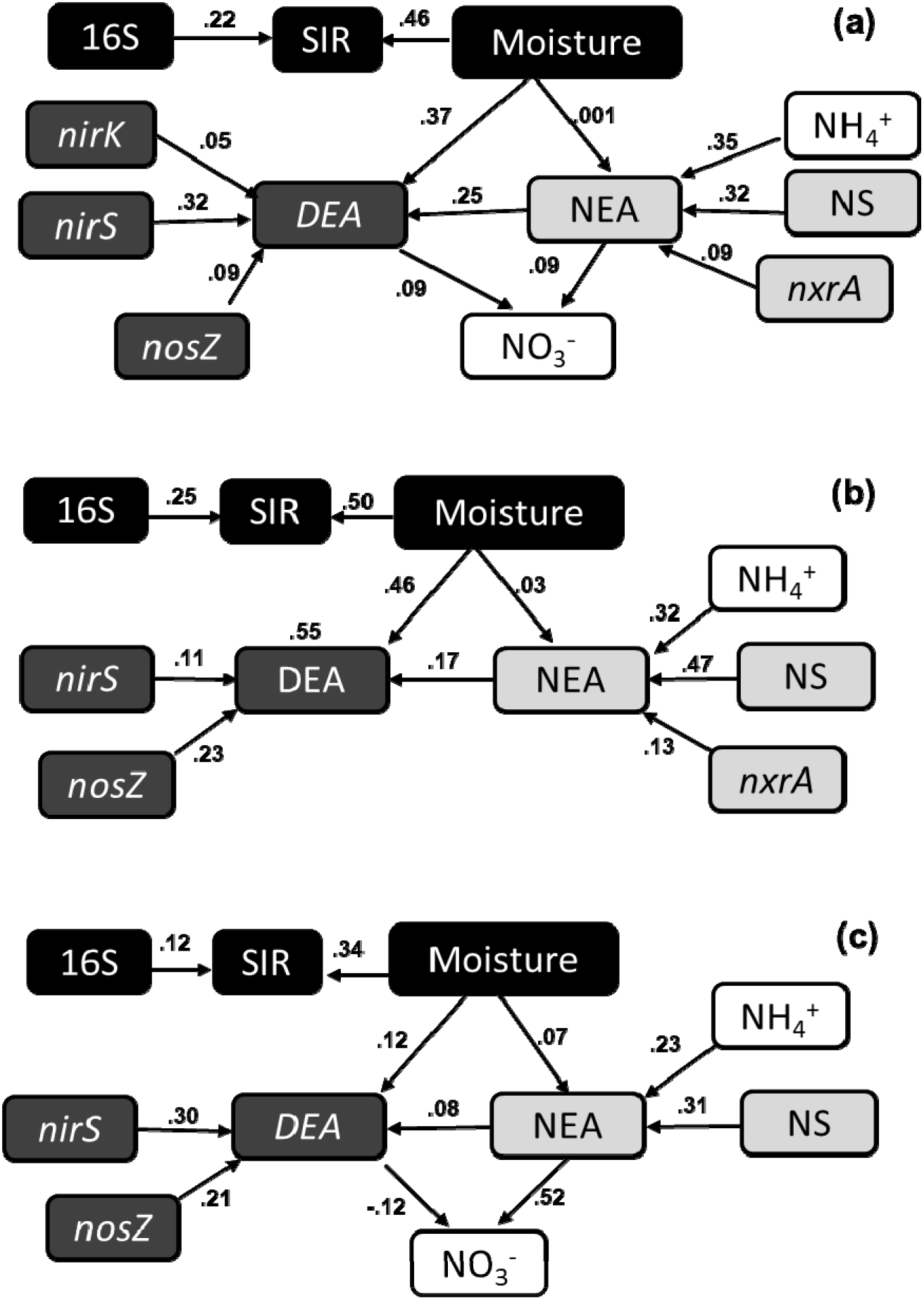
Structural equation models (SEM) bridging the effects of microbial and environmental factors. SEMs including the Substrate induced respiration MicroResp (SIR), nitrification (NEA), denitrification (DEA), gene community abundances (16S *rRNA*, *nxrA*, *NS*, *nirK*, *nirS*, *nosZ*) and environmental factors (soil moisture, NH_4_, NO_3_) in **a, b, c** are built using the data from treatments; SR, D, SR+D, respectively. The values shown beside the arrows are the path coefficients and correspond to the standardized coefficients calculated based on the analysis of correlation matrices. Values only for the significant relations (that fit the model, P > 0.05) are indicated in the SEMs.

## 4. Discussion

This work reported the resilience including the resistance and recovery capabilities at subalpine grassland ecosystems. Investigating the impact of applied climate change treatments was crucial since these ecosystems are under drastic influence of global climatic changes which may ultimately disrupt their functioning and the linked ecosystem services (Berauer et al., 2019; Bernard et al., 2019; Schirpke et al., 2013; Wipf and Rixen, 2010; Vittoz et al., 2009). Our results demonstrated the impact of successive droughts and early snowmelt climatic extremes on subalpine grassland ecosystem functioning including nitrification and denitrification as model processes and such findings give valuable insights into overall N cycling resilience (Chen et al., 2020; Dai et al., 2020).

The quantification of resistance and recovery by comparing the relative change in D, SR, and SR+D treatments over time compared to control showed the strong impact of different climate change treatments on microbial activities and the corresponding functional groups. Cluster analyses approach used to determine the effect of treatments within each block at different sampling times revealed five different clusters. In each cluster, specific treatments grouped separately to show that effect of one specific treatment does not influence any block. The further effect of outliers was analyzed to know the difference in descriptive values after non-consideration of outliers. Overall, broad scale function – SIR and the size of total bacterial community - showed low resistance but high recovery for all treatments. The finding strengthened the fact that this function is carried out by a wide range of microbial communities which generally exhibit a fast recovery (Ingrisch et al., 2020). SIR represents the broad scale microbial activity hence disturbing a part of the community could not affect the overall function on the long term. Yet, short term reduction in SIR upon droughts can be associated to lower microbial activities induced by water deficit conditions (Bernard et al., 2019; Bloor and Bardgett, 2012). Moreover, broad scale functions are reported to be less sensitive to applied disturbance (Chaer et al., 2009). Like respiration, decomposition processes are generally driven by broad scale communities and weak impact of climatic stresses for these processes have been reported in a joint experiment (Bernard et al., 2019), while high recovery of microbial biomass to climatic stress have been observed in other mountainous grasslands (Piton et al., 2020). These observations are also in accordance with our assumptions on functional redundancy concepts (Jia and Whalen, 2020; Griffiths and Philippot, 2013).

The impacts of extreme climatic events including droughts on microbial N cycling are less clear (Fuchslueger et al., 2019). Climate change treatments showed contrasting impacts on N cycling processes with denitrification (community gene abundance and activity) showing less resistance to drought (D) as compared to nitrification (both for community gene abundances and activities). For example, at T3 where snow removal and droughts occurred in the same mesocosms (SR+D), DEA was first strongly reduced followed by an incomplete recovery. Such observations were in contrast to NEA which was affected by snow removal (SR) but recovered completly at T4. Our findings confirmed a strong influence of droughts and associated soil moisture deficits specifically for DEA (Di et al., 2020; Philippot et al., 2009). This adverse influence of applied stresses was pronounced when both climatic stresses (SR+D) occurred indicating the additive effect of SR with declined nitrification upon earlier snowmelt. Interestingly, the observations were both stress and community specific, with variable response for microbial community size and functions. The collapse in microbial pool size upon snowmelt was observed while studying the niche differentiation perspectives in soils (Sorensen et al., 2020). In nearby subalpine grasslands, a strong turnover by nitrifying and denitrifying microbial communities was previously shown upon natural variations in snowpack depth (Jusselme et al., 2016). In wake of climatic extremes and considering the functional stability scenarios, such multiple cascading effects are important to consider not only from microbial community perspectives but also for related ecosystem services. For example, these cascading effects can be crucial for N-related microbial activities which can adversely impact greenhouse gas emissions (i.e., N_2_O in particular), soil fertility and plant productivity, hence ask for adaptive management strategies to counter the unforeseen consequences (Griffis et al., 2014; Di et al., 2014; Bouwman et al., 2009).

The reduction in gene abundances of nitrifiers and denitrifiers upon drought (D) or snow removal (SR) delineated the strong impact of climate change treatments in studied grasslands. Denitrifiers are usually reported as more tolerant to droughts (Hammerl et al., 2019) than nitrifiers (Austin and Strauss, 2010). Here, in line with DEA, denitrifiers’ abundances were also negatively impacted by droughts, especially after the second one. Like DEA, *nirK* denitrifiers’ abundances were severely lowered and could not show a complete recovery at T4. DEA seems to be related to various denitrifying communities including *nirK*. In agreement, DEA and abundance of *nirK* denitrifiers have been found to be correlated upon variable snow depths under similar subalpine grasslands (Jusselme et al., 2016). Conversely, Wertz et al. (2007) found that the abundances of denitrifiers were less affected by heat stress in an agricultural soil, and this was associated to high functional redundancy.

Soil abiotic properties are crucial in estimating soil N cycling processes, and resilience of microbial communities upon applied stresses may be influenced by inherent soil edaphic characteristics (Berard et al., 2015). In our study, although we found strong positive correlations of DEA with %C, %N, DON, SOM, soil moisture, NEA, and denitrifiers’ abundances, interestingly these factors shifted during the studied time. For example, when grasslands were exposed to combined snow removal and drought (SR+D), soil moisture along with *nirK*, *nirS* and *nosZ* were correlated to DEA (r = 0.33, 0.15, 0.32, 0.38; p < 0.005). While soil moisture and *nirK* were not correlated to DEA (r = 0.04; p > 0.64) in the CTRL. These observations showed the variable dominance of different denitrifiers in D, SR and CTRL treatments. Similarly, DEA has been associated differently to these variables across a range of environments (Chen et al., 2020; Chen et al., 2019; Petersen et al., 2012; Chroňáková et al., 2009, Dong et al., 2009). At T4, DEA could not recover demonstrating a severe impact caused both by drought (D) and the additive effect with snow removal (SR+D). Moreover, the impact on nitrification during SR significantly passed on to DEA which collapsed upon the second drought showing the cascading effect for SR+D.

These pronounced impacts on DEA and corresponding denitrifiers’ abundances were also explained through SEM (Fig.5). Possibly the system, (as described by Schnellnhuber et al. (2006)) upon disturbance was ‘pulsed’ (respond on alteration but return) and ‘pressed’ (respond on sustained alteration so shift) where ‘press’ brings the ecosystems to a tipping point (as observed for DEA in our study). This may shift the system to an irreversible new state (Wall, 2007), or at least may lead to altered recovery patterns (Seabloom et al., 2020). Furthermore, such changes in denitrifiers’ relative abundances may also result in selection of certain stable/unstable groups. Such extreme climatic events may consequently impact (increase) N losses through NO_3_ leaching or N_2_O emissions (de Vries and Shade 2013). In addition, studies in other ecosystems have suggested that these functional guilds may have contrasting resilience abilities against various disturbances (Capdeville et al., 2019).

Though SIR was related to soil moisture and *16S rRNA* when SEM was fitted, NEA was highly linked to both soil NH_4_^+^ and *NS* nitrifiers’ abundance hence showed a weak effect of soil moisture on NEA. Therefore, *NS* communities might have played the role as dominant nitrifiers in these grasslands. Higher abundance of *NS* nitrifiers than *nxrA* under drought as well as the dominant role of *NS* nitrifiers in these soils was earlier shown in other soils with low activity (Attard et al., 2010). However, here DEA was related both to soil moisture and abundances of *nirS* denitrifiers throughout the experiment except at T3 when it was *nosZ*, suggesting community selection or adaptation is an important phenomenon under changing climate or stress conditions. After the second drought at T3, a cascading effect of snow removal (SR) might have shifted the community dynamics from *nirS* to *nosZ* with better adaptation of the *nosZ* denitrifiers. The denitrifiers responded differentially, *e.g.,* in contrast to *nirK*; *nirS* and *nosZ* abundances recovered, showing a community specific impact (Karlowsky et al., 2018) of drought and snow removal. The *nirK* denitrifiers were reported as less abundant (Peterson et al., 2012), and relatively more sensitive to environmental change than *nirS* denitrifiers in subalpine and agricultural soils (Szukics et al., 2019; Hallin et al., 2009). Though the findings implied by Dai et al. (2020) contrasted with these observations, seasonally driven drought effects have been reported for *nirK*, *nirS* and *nosZ* abundances in grassland ecosystems (Hammerl et al., 2019). Moreover, the denitrifiers harboring *nirK* genes might not contain *nosZ* genes, in agreement to the discovery that not all denitrifiers contain *nosZ* (Hallin et al., 2018), which may ultimately disturb the overall nitrous oxide budget from studied soils.

We did not find any significant effect of plant functional composition and fertilization on the investigated soil microbial parameters, suggesting they were not the main drivers of the studied processes in our experimental set-up. Consequently, the trait-centered influence of abiotic variables on broad and N-related microbial abundances and activities, in response to climatic stresses, was not observed here but, instead, their direct effect (Griffin-Nolan et al., 2019). Yet, leaf or root specific traits, rather than plant community composition, might have affected the nutrient availability and ultimately associated microbial processes. This could explain why we found that soil nutrient contents along with community abundances were driving the N cycling processes in our study. Interestingly in similar grasslands, no direct effect of fertilization was observed but only specific plant traits were affecting the same microbial activities (Legay et al., 2016). This suggests a competition for nutrients between plants and microbes, as observed following snowmelt in nearby subalpine grasslands (Legay et al. 2013), resulting here in a non-significant fertilization effect on microbial parameters during the study period. However, such management strategies including fertilization practices may favor certain microbial communities ultimately causing fast resilience to stress conditions (Piton et al., 2020). It is noteworthy that due to the climatic extremes, the responses showed by the various nitrifiers’ and denitrifiers’ groups were different to CTRL indicating an unanticipated internal competition for initially available nutrients. Importance of such competitions will be crucial to further investigate, especially in ecosystems facing climatic extremes, since these are suggested to be critical in controlling ecosystem N cycling (Legay et al., 2013; Grigulis et al., 2013, Le Roux et al., 2013).

## 5. Conclusions

This work clearly demonstrated a strong and unprecedented impact of climate extremes such as early snowmelt and recurrent droughts, alone or in combination, on soil microbial communities both at broad (i.e., soil respiration) and N-related (nitrifiers’ and denitrifiers’ activities and abundances) scales. Droughts showed a low and short-term impact on bacterial total respiration supporting their hypothesized high resilience (resistance and ability to recover). Though nitrification activity and community gene abundances of corresponding functional guilds exhibited relatively strong resistance to summer droughts, they declined in response to early snowmelt leaving strong impact on denitrification. Our findings inferred that predicted climatic extremes may carry severe consequences for soil microbial community functioning in subalpine grasslands ultimately resulting in shifts in N availability, plant community diversity, and associated ecosystem services.

## Acknowledgements

This work was part of the ERA-Net BiodivERsA project REGARDS (ANR-12-EBID-004-01). Quantitative PCRs were carried out at the platform DTAMB (IFR 41, Université Lyon 1) and N cycling activities at the platform AME (UMR5557-UMR1418). We want to thank all the people who helped us in the field or in the lab: C. Arnoldi, A. Foulquier, N. Guillaumaud. This research was conducted at ‘Jardin du Lautaret’ (UMS 3370 Univ. Grenoble Alpes – CNRS), a member of the DIPEE Grenoble-Chambery, the FREE-Alpes Federation (FR n°2001-CNRS), and of the AnaEE-France (ANR-11-INBS-0001AnaEE-Services). This research was conducted within the Long-Term Socio-Ecological Research (LTSER) platform Zone Atelier Alpes, a member of the eLTER network.

## Supplementary Figures

**Fig. S1.**
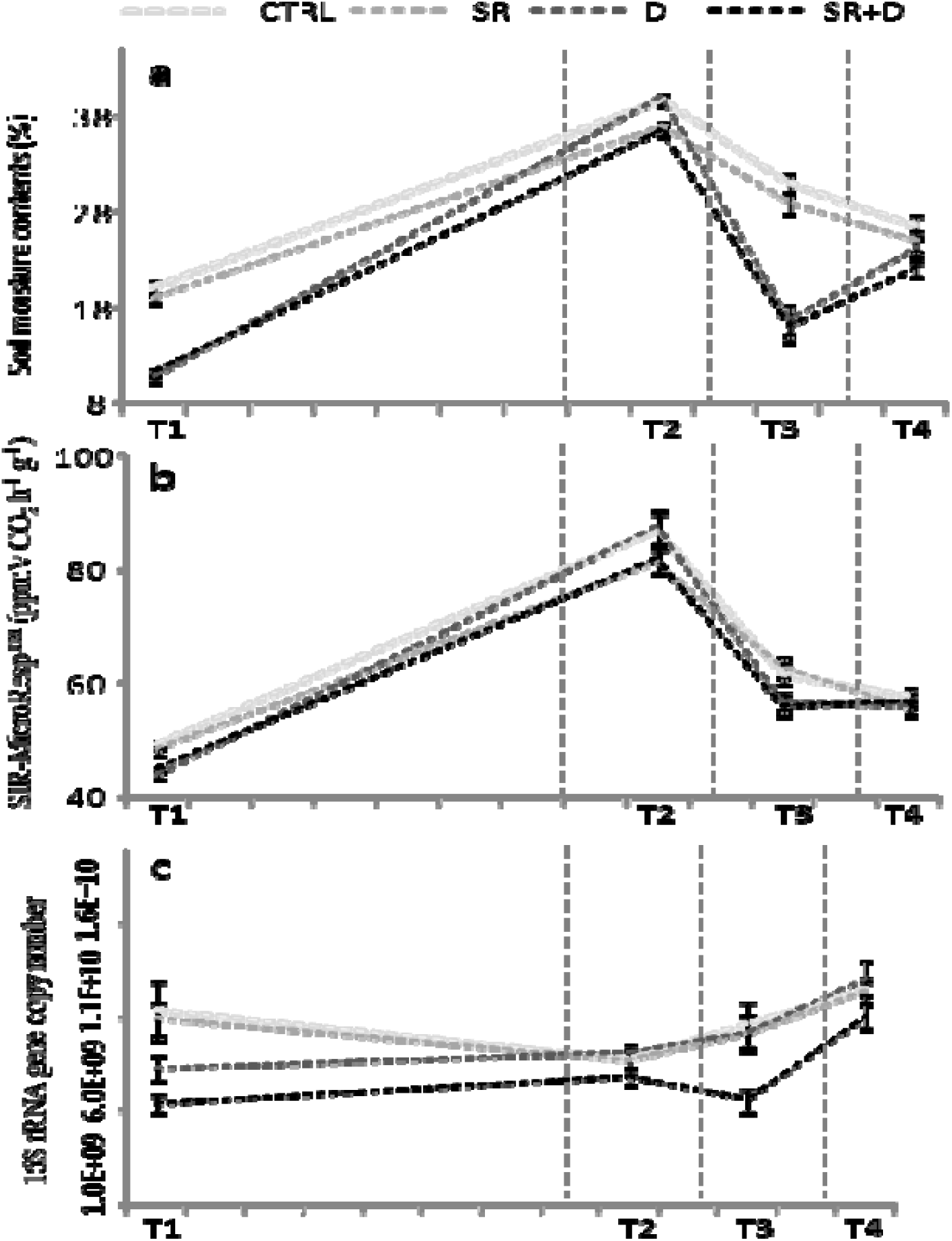
**a)** Soil moisture in percentage of dry soil; **b)** SIR-MicroResp™ (ppmV CO_2_ h^−1^ g^−1^); **c)** Abundance of total bacterial community (number of 16S *rRNA* gene copies g^−1^ dry soil).

**Fig. S2.**
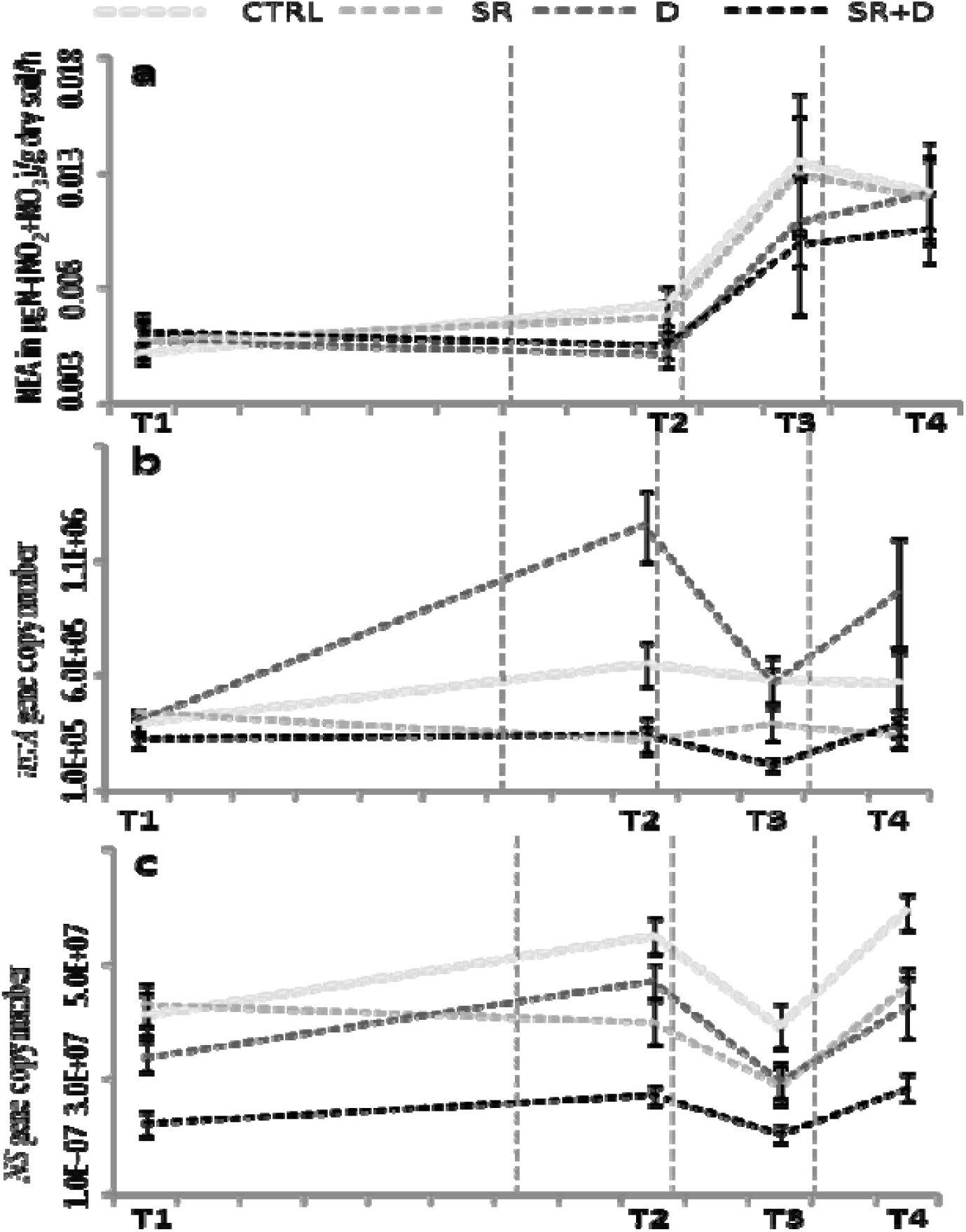
**a)** Nitrification enzyme activities quantified in μgN-(NO_2_+NO_3_)/g dry soil/h for different treatments at various sampling dates; **b)** Abundance of *nxrA* nitrite reductase (number of *nxrA* gene copies g^−1^ dry soil) for different treatments at various sampling dates; **c)** Abundance of *NS* nitrite reductase (number of *NS* gene copies g^−1^ dry soil) for different treatments at various sampling dates.

**Fig. S3.**
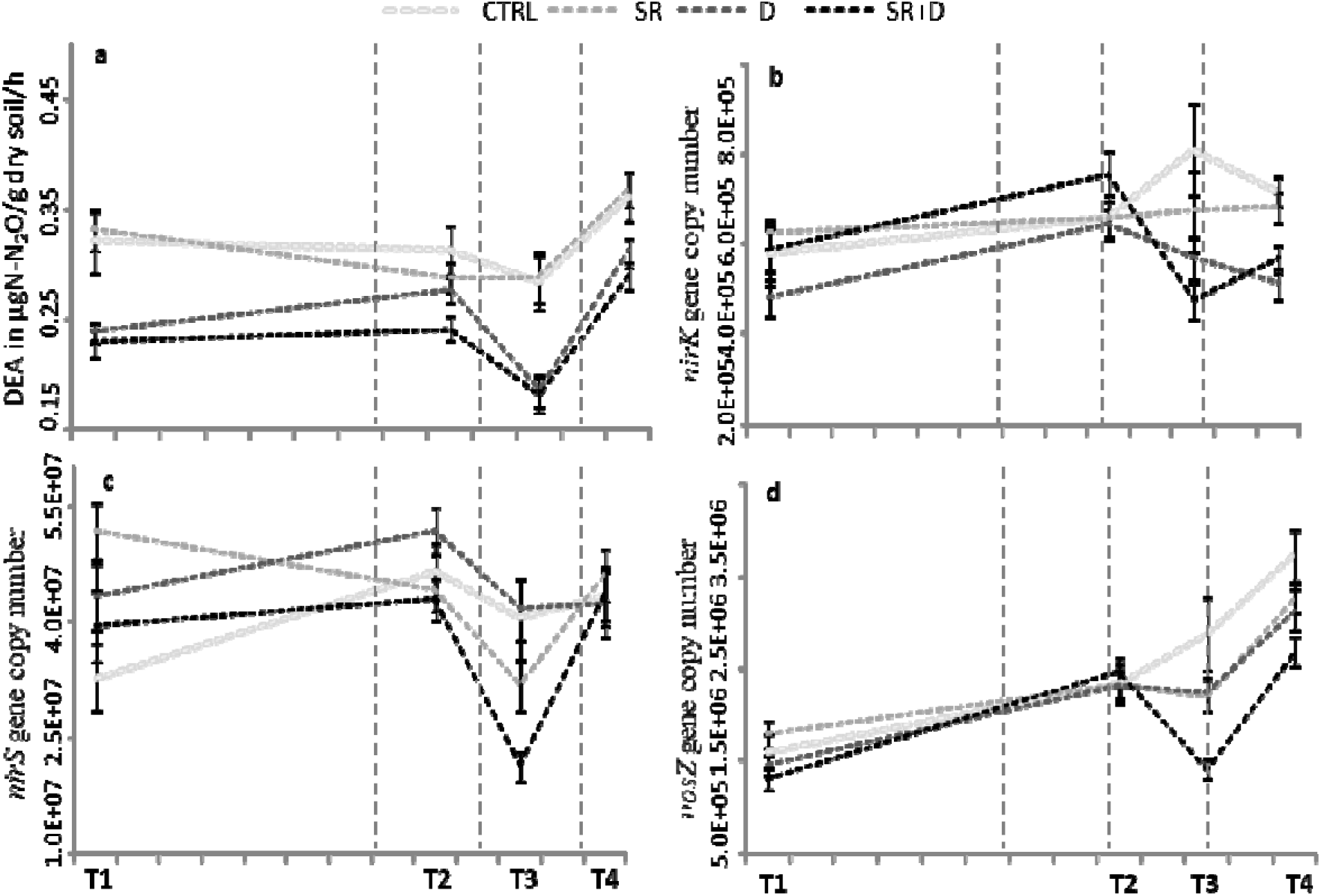
**a)** Denitrification enzyme activities determined in μg N-N_2_O/g dry soil/h for different treatments at various sampling dates; **b)** Abundance of *nirK* denitrifiers (number of *nirK* gene copies g^−1^ dry soil); **c)** Abundance of *nirS* denitrifiers (number of *nirS* gene copies g^−1^ dry soil); **d)** Abundance of *nosZ* denitrifiers (number of *nosZ* gene copies g^−1^ dry soil) for different treatments at various sampling dates.

